# Rediscovery and ecology of the Capulin Mountain Alberta arctic butterfly: informing data collection for rare endemics

**DOI:** 10.1101/2025.10.23.684241

**Authors:** Simon M. Doneski, Quinlyn Baine, Kelly Ricks, Steven J. Cary, Kelly B. Miller

## Abstract

Insect conservation in the southwestern United States is hindered by limited information on distribution, phenology, and habitat, especially for small-range endemics.

The Capulin Mountain Alberta Arctic butterfly (*Oeneis alberta capulinensis* F.M. Brown, 1970), a New Mexico Species of Greatest Conservation Need, is restricted to the Raton Mesa Complex and threatened by isolation, low dispersal, and extirpation from its type locality.

We surveyed historic and potential sites to document current distribution, refine flight phenology, investigate larval host plant use, and assess habitat condition. Our results confirm extant populations, identify the larval host plant (*Festuca idahoensis* Elmer), and describe habitat as montane grasslands with limited woody encroachment.

A habitat suitability model predicted additional potentially occupied areas, highlighting under-explored habitat for future monitoring.

This study provides the first comprehensive ecological assessment of *O. a. capulinensis* and establishes a framework for documenting poorly known endemic insects to guide future research and survey design.

## Introduction

Effective conservation action requires a background of data on population dynamics, distribution, ecology and basic natural history that is lacking for the majority of insects, particularly in the southwestern region of the United States (Cardoso *et al*., 2011). This region is characterized by diverse landscapes from the arid to the alpine that foster high species richness and morphological diversity (Danks, 1994; Stein, 2002). Habitat heterogeneity in the state of New Mexico, home to ecological diversity ranging from the Chihuahuan Desert to the Southern Rocky Mountains results in not only high diversity of species but also high rates of local endemism. Many species have evolved characteristics in biogeographic “islands” that keep them intrinsically tied to a specific habitat in a specific geographical location (Gottfried *et al*., 2005). Endemic taxa are often restricted to narrow niches and small populations leading to reduced genetic diversity and limited adaptive potential when confronted with environmental change, thereby increasing extinction risk (Chichorro *et al*., 2019; Staude *et al*., 2020; Manes *et al.,* 2021). To protect endemics, especially those with particularly small ranges, conservationists should undertake specific targeted efforts to collect necessary data for adequately assessing conservation risks for populations.

One such small-range endemic is the Capulin Mountain Alberta arctic butterfly (*Oeneis alberta capulinensis* F.M. Brown, 1970), herein referred to as the Capulin arctic. The Capulin arctic is one of the southernmost subspecies of the Alberta arctic a cold adapted species which is found across Canada and the northern Rocky Mountains (Cary & Toliver, 2025). During the last glacial maximum the Alberta arctic extended its distribution all the way south to New Mexico in the period of warming since then most Alberta arctics have returned to more northern latitudes but several isolated populations remain in New Mexico. The Capulin arctic is an isolated alpine grassland specialist butterfly that is known only from a single complex of elevated mesas and volcanic peaks on the east flank of the Southern Rocky Mountains along the New Mexico and Colorado border known as the Raton Mesa Complex (Johnson *et al.,* 2004; Cary & Toliver, 2025). Subspecies like this, with extremely restricted ranges and seemingly limited habitat requirements, are highly susceptible to threats and may easily become extinct within our lifetimes without our assistance. However, knowledge about its natural history is limited. Given recent increase in interest in arthropod conservation in New Mexico, we have a unique opportunity to gather essential data on rarely observed and potentially-threatened endemics such as the Capulin arctic.

The years of 2023 and 2024 marked significant progress for conservation of insects in New Mexico. In January of 2023, the state’s first protected insect, the Sacramento Mountains Checkerspot Butterfly (*Euphydryas anicia cloudcrofti* Ferris and R. Holland, 1980) was federally listed as Endangered (U.S. Fish and Wildlife Service, 2023). Soon after, the state got its second federally-protected insect with the Threatened listing of the Nokomis Silverspot Butterfly (*Argynnis nokomis nokomis* [Edwards, 1862]) in February 2024 (U.S. Fish and Wildlife Service, 2024). This laid the groundwork for considerable change in insect conservation in the state.

Recently, the New Mexico Rare Arthropods Resource was developed to identify and track arthropods of conservation concern (New Mexico Rare Arthropods Technical Council, 2025). Using and expanding on this resource, the New Mexico Department of Game and Fish included pollinating insects as Species of Greatest Conservation Need (SGCN) for the first time in twenty years in the 2025 State Wildlife Action Plan revision. Ninety-three insects were designated as SGCN, including the Capulin arctic (New Mexico Department of Game and Fish, 2025). Then, in early 2025, the authority of the New Mexico Department of Game and Fish was expanded to protect non-game wildlife species in need of conservation and the definition of wildlife in the state’s constitution was amended to include all invertebrates, including butterflies (New Mexico Legislature, 2025).

The 2025 field season was the first in New Mexico where butterflies were recognized as valid “wildlife” for conservation attention. The Capulin arctic is one of the first SGCN butterflies to eclose as adults in spring with males starting to fly in early May and females emerging shortly afterwards (Cary & Toliver, 2025). Extreme flight dates for the Capulin arctic in New Mexico range from 3 May–5 Jun with peak population size in mid-May (Cary & Toliver, 2025). This butterfly subspecies occurs only in the Raton Mesa Complex a series of high-elevation mesa tops and volcanic peaks straddling the northeast border New Mexico and southeast border of Colorado. The Raton Mesa Complex includes windswept fescue grassland, the general habitat of the subspecies.

One of the concerns about the conservation status of the Capulin arctic originates from Brown’s (1970) original description, in which he remarks that he believes the subspecies to be experiencing high levels of inbreeding depression (Brown, 1970). He supported this with his observation that the Capulin arctic rarely flies and in all his time working with the butterfly he only observed females walking on the ground, potentially pointing to an extremely low dispersion rate for this butterfly (Brown, 1970). Further survey work completed in 2003 and 2004 increased concern for the Capulin arctic when it was discovered that the population was evidently extirpated from its type locality, Capulin Volcano National Monument in the eastern region of the Raton Mesa Complex (Johnson *et al.,* 2004). The loss of an entire population from a well-protected area highlights the precarity of small-range endemics in general. It also suggests a need for regular monitoring of extant Capulin arctic populations to ensure their survival or persistence.

Our main goal in this study was to expand knowledge of the Capulin arctic in New Mexico. We investigated its distribution, phenology (especially flight phenology), life history, and habitat. The only known larval host plant for the Alberta arctic is a bunchgrass in the genus *Festuca* (Cary & Toliver, 2025; New Mexico Rare Arthropods Technical Council, 2025).

Abundance of this grass is evidentially essential to the persistence of this butterfly. Our goal was to determine if that is the case still for the Capulin arctic and to identify the specific *Festuca* species involved and document its role as a host plant. We also aimed to assess the condition of the subspecies’ habitat in light of recent drought conditions across New Mexico. Finally, we used natural history information we collected in the field to inform a model of suitable habitat constructed from remote sensing data and to predict where additional extant populations may remain.

## Materials and Methods

### Survey

Our study incorporated the publicly accessible grasslands that occur above the tree line east of I-25 in Colfax and Union counties in New Mexico (Fig. 1). These grasslands are associated with volcanic summits and basalt-capped mesas from Bartlett Mesa on the western margin and eastward to Sierra Grande, a massive and extinct shield volcano (Johnson *et al*., 2004).

**Figure 1.**
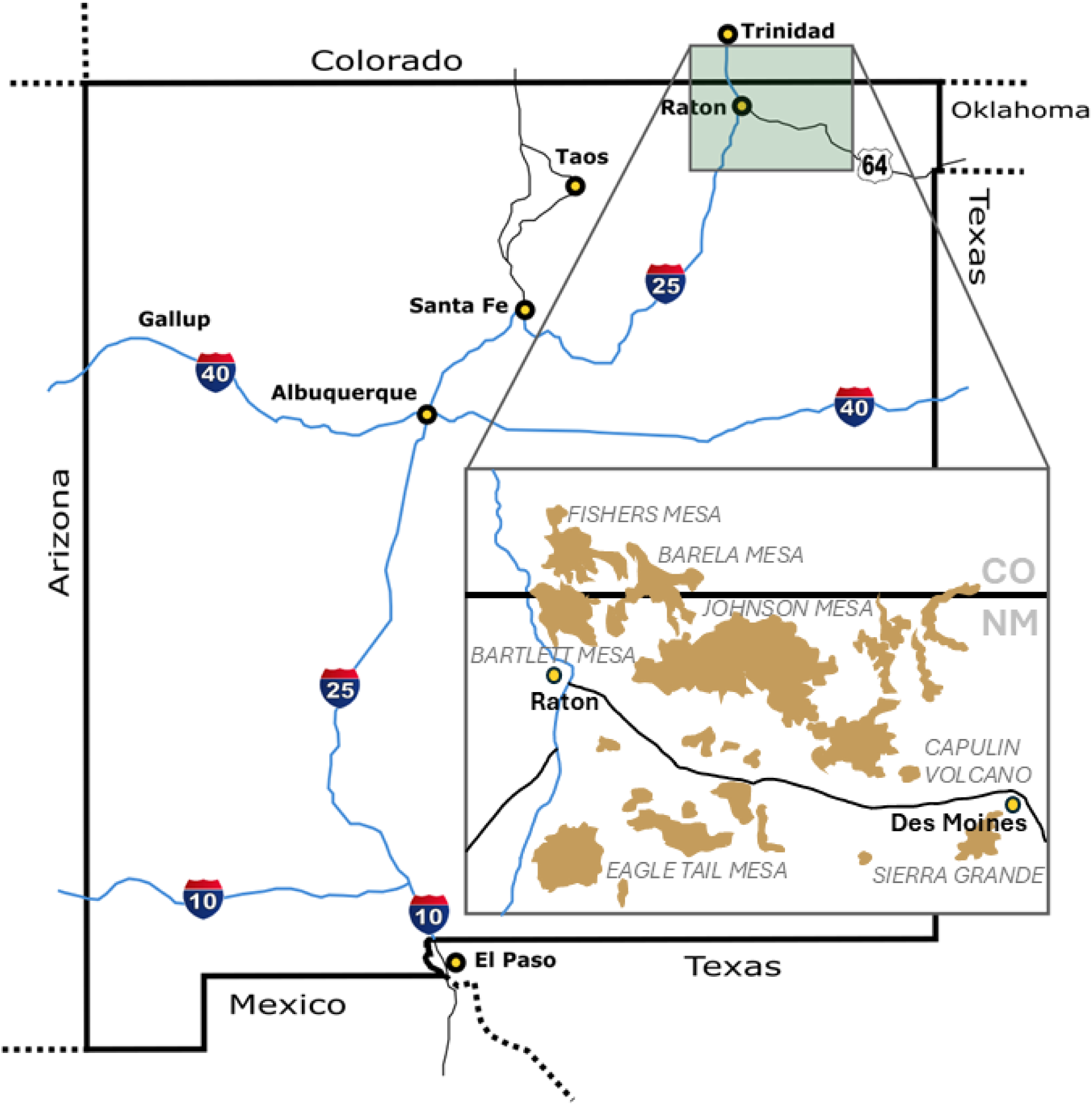
Map of survey area and the Raton Mesa Complex.

Much of this region is privately owned and not publicly accessible. Collections were made on New Mexico State Public Lands. We also accessed Sugarite Canyon State Park and Capulin Volcano National Monument (the type locality for the Capulin arctic).

We investigated previously documented populations and attempted discovery of new ones. At these study sites direct visual observation and binoculars were used to locate adults in flight and used aerial insect nets to capture adults. A total of six individuals were collected as voucher specimens. These individuals were deposited in the Museum of Southwestern Biology’s Division of Arthropods. (MSBA, K.B. Miller, curator; MSBA98995, MSBA99002, MSBA99012, MSBA99023, MSBA99026, and MSBA99030). Additionally, tissue was collected and preserved in the Museum of Southwestern Biology’s Division of Genomic Resources (MSB DGR, M. Anderson, curator; AAN5W, AAN5Z, AAN60, AAN5X, AAN5P, AAN5Y).

Specimens were identified using Cary & Toliver (2025), and expert opinion from SJC, and the taxon concept used was the original description (Brown, 1970). Cameras were used to capture images of adults observed on the ground when possible (Fig. 2A).

**Figure 2.**
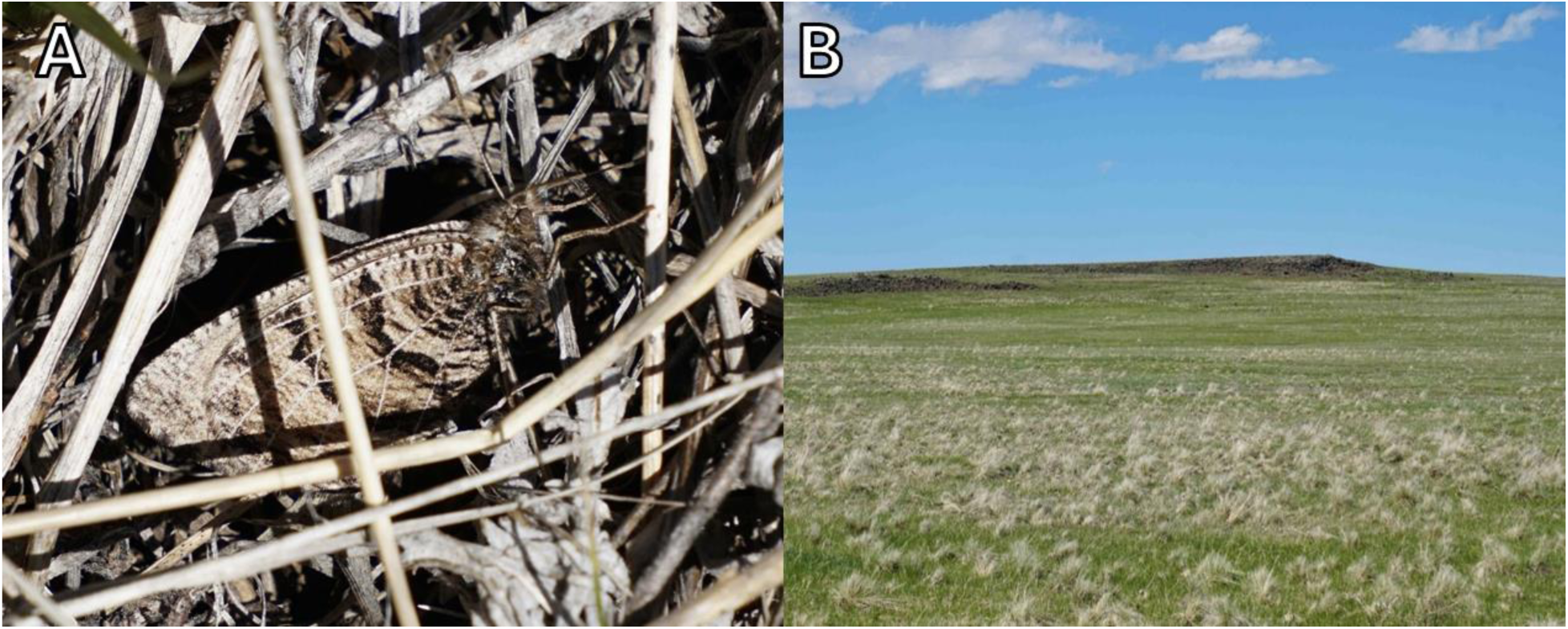
A. Adult Capulin arctic resting in grassy debris at base of Festuca idahoensis.B. Johnson Mesa site, pale, dry bunches of Festuca idahoensis in foreground. Colfax Co., NM; May 15, 2025; photos by Kelly Ricks.

In each area searched, the status and abundance of *Festuca* species was evaluated (Fig. 2B). At each site a random route was taken through the visually identifiable stands of suitable habitat. Vouchers of the host grass were collected from Johnson Mesa and Bartlett Mesa to confirm identification in the lab. This was done during surveys and once in August for reproductive morphology,

### Habitat Modeling

After completing surveys, it became clear that the *Festuca* host grass was present throughout the area in scattered but distinct patches. These were fairly evenly dense and easy to recognize from a distance because of the bunch-grass shape and tall tufts of dead prior-year vegetation that gave plants a light, faded color compared to other lower-profile and greening-up grasses (Fig. 2B). Based on this observation, a new method was developed to detect potentially suitable Capulin arctic habitat, an aerial imagery examination of grasslands in the Raton Mesa Complex delimited by elevation to identify patches the color of *Festuca*.

For the habitat suitability model, Landsat Operational Land Imager (OLI) 30 meter resolution cloudless images for Colfax and Union Counties taken in May-July 2014 were used. A layer was created from rasters of 10 meter resolution Digital Elevation Models (DEM) prepared from images taken in 2005 by the United States Geographical Survey. Both datasets were provided by the New Mexico Resource Geographic Information System (NM RGIS) managed by the University of New Mexico’s Earth Data Analysis Center.

A maximum likelihood classifier (MLC) in ArcGIS Pro 3.4.0 (Esri) was trained to categorize land cover classes from the aerial imagery using a supervised approach. Three classes of land cover were designated and subsequently confirmed by field observations. These were: 1. forest, characterized by dense trees, 2. unsuitable grassland, characterized by rhizomatous, non- bunching grasses, and 3. suitable grassland, where Capulin arctic has been observed, characterized by *Festuca* bunchgrasses. The samples used for training were 16 digitized polygons (between 0.02-19 km^2^) that defined the 3 classes, from which the algorithm computed statistical class signatures under the assumption of a multivariate normal distribution. To improve class separability, we incorporated segment attributes, including mean digital number and converged color, as predictor variables. The classifier then calculated the posterior probability for each 30 m segment across all classes and assigned each to the class with the highest likelihood.

From the classified segments, the class “suitable grassland” was isolated and then the layer was clipped to two options for minimum elevation: 1. 2,475 m from the lowest known occurrences of Capulin arctic, and 2,300 m as a generous buffer to account for uncertainty of habitat requirements. Total area of both models was calculated to represent the total amount of suitable habitat that exists in New Mexico.

However, based on well-supported evidence that Capulin arctic is now extirpated from the top of Capulin Mountain, which is an extremely small area compared to nearby mountains and mesas where it persists (0.8 km^2^ total area above 2,300 m in elevation), we estimated that sufficient size of habitat is also important for the butterfly’s survival. Assuming that the area of Capulin Mountain is below a hypothetical area threshold of habitat suitability, a minimum habitat requirement of 0.8 km^2^ was assigned and the models were trimmed to include only areas of continuous habitat above that size threshold to more accurately represent where Capulin arctic may be extant today.

Finally, to get an estimate of suitable habitat that is on non-privately-owned land, as a way to gain an understanding of accessibility and protected status of these sites, the models were clipped to a Bureau of Land Management New Mexico Surface Ownership map (sourced from NM RGIS, last updated in 2022), and total area of suitable habitat under public ownership was calculated.

## Results

### Survey

Meaningful field observations were acquired at several locations during the 2025 flight season. At Johnson Mesa (Sec. 16, T31N, R25E) on the mornings of 13–14 May (36.921948, -104.287003), we conducted two to three hours of sampling and found abundant *Festuca*, later identified as *Festuca idahoensis* Elmer, along with high numbers of Capulin arctics. Over the two days, an estimated total of 70–80 adults were observed, including both males and females, most in very good condition. Two specimens (one male and one female) were collected as vouchers (catalog numbers MSBA98995 and MSBA99023). A second site at Johnson Mesa (Sec. 16, T31N, R26E) was surveyed in the afternoons of 13–15 May (36.92217, -104.1783), following different routes each day primarily south and southwest of Dale Mountain (Fig. 2B). The presence of *F. idahoensis* and Capulin arctics was similar to Site 1, although the total number of adults observed was lower, approximately 35–40 individuals. Three voucher specimens were collected (catalog numbers MSBA99002, MSBA99030, MSBA99012).

Horse Mesa (SE1/4 of Sec. 36, T32N, R24E) was surveyed on 14 May in the afternoon (36.965426, -104.340299). This steep and rugged east side-slope is largely wooded and did not provide habitat for the Capulin arctic. Nevertheless, this area may support other New Mexico rare butterflies, including *Lon hobomok wetona* (Scott, 1981) and *Argynnis hesperis ratonensis* (Scott, 1981). Capulin arctics were previously recorded along the west rim as recently as 20 May 2008, but no observations have been made since despite several survey attempts.

On 15 May, we surveyed the utility service road to the summit of Sierra Grande (36.70560, -103.87681). Large amounts of *F. idahoensis* were present near the summit and extending down the steep northeast slope. Despite the strong winds and resulting windchill, the fescue-filled lee slopes offered suitable habitat; however, no arctics were encountered.

Bartlett Mesa, at the northeastern edge, was surveyed on 16 May in the morning (36.98341, -104.40572). Conditions were cool and windy, and few butterflies were active even after 11:00 AM. One Capulin arctic was confirmed with an additional probable sighting. The mesa top contains prairie habitat with variable concentrations of fescue, and one voucher specimen was collected (MSBA99026). Little Horse Mesa, Sugarite Canyon State Park (36.98252, -104.40119), was also surveyed on 16 May in the morning, where few butterflies of any kind were observed.

Routine monitoring at Capulin Volcano National Monument (CAVO; 36.78313, - 103.96778) began in April and continued throughout the flight period, focused on areas still containing fescue grasses, including the type locality. The Capulin arctic was not observed at CAVO during the 2025 season.

### Habitat Modeling

Four models were produced to estimate the extent of suitable habitat for Capulin arctic in New Mexico. In the most liberal models, which did not incorporate a minimum habitat size, 155.9 km² of suitable habitat was predicted at a minimum elevation of 2,300 m, and 120.5 km² at a minimum elevation of 2,475 m (Fig. 3). When filtered using a minimum habitat size of >0.8 km², predicted habitat area was reduced to 124.8 km² at 2,300 m elevation and 113.7 km² at 2,475 m elevation. An estimated range of 113.7–155.9 km² (70.6–96.9 mi²) of potential Capulin arctic habitat in New Mexico is estimated (Fig. 3). In the most conservative model, suitable habitat was predicted in seven named geographic regions of New Mexico: Sierra Grande (Fig. 4B), Johnson Mesa, Little Mesa, Barela Mesa, Horse Mesa, Little Horse Mesa, and Barlett Mesa (Fig. 4A).

**Figure 3.**
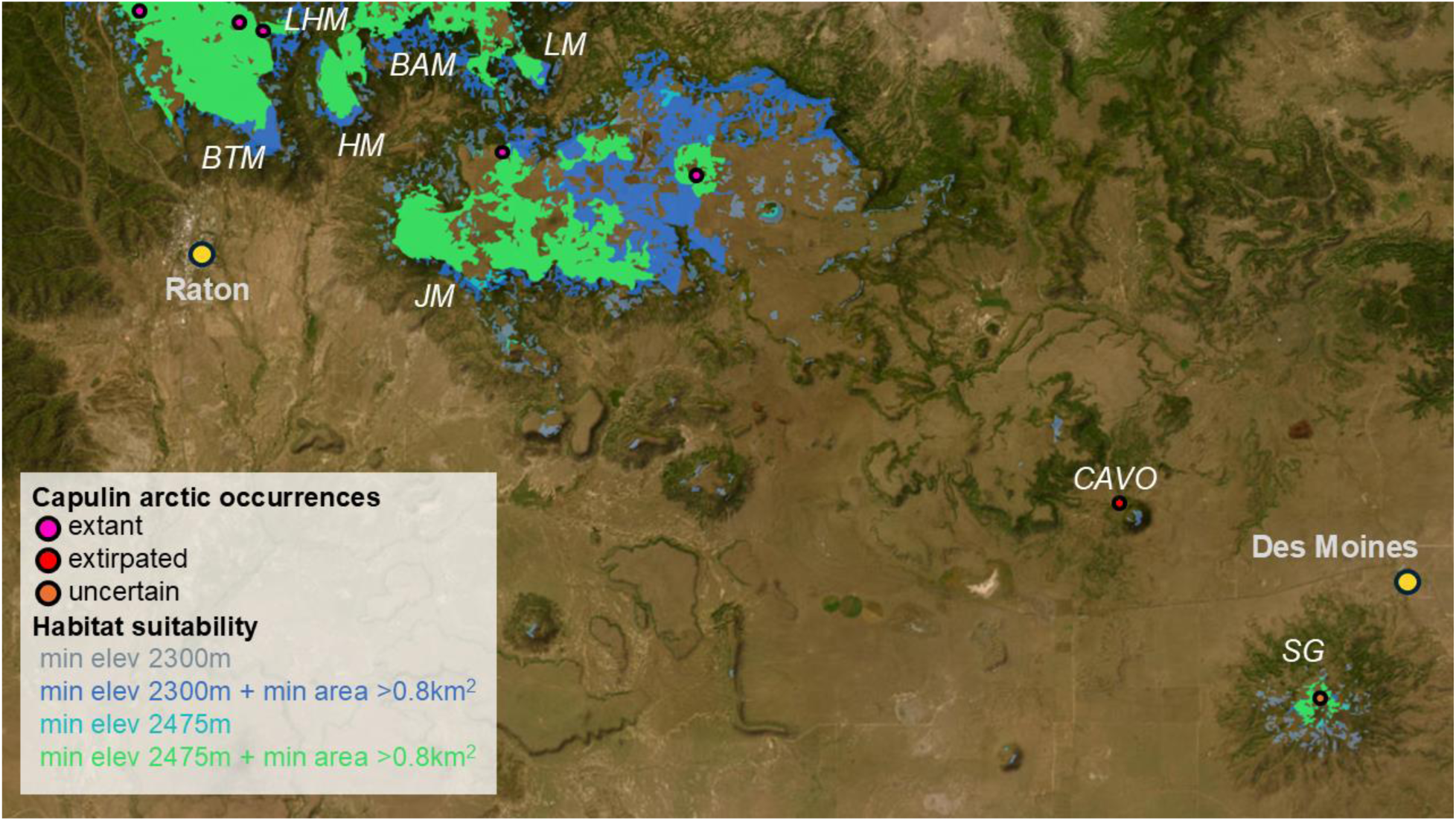
Capulin arctic occurrences and results of habitat suitability modelling using remote sensing data. Basemap imagery by Esri et al. (2025). BTM = Barlett Mesa, HM = Horse Mesa, LHM = Little Horse Mesa, BAM = Barela Mesa, LM = Little Mesa, JM = Johnson Mesa, CAVO = Capulin Volcano National Monument, SG = Sierra Grande.

**Figure 4.**
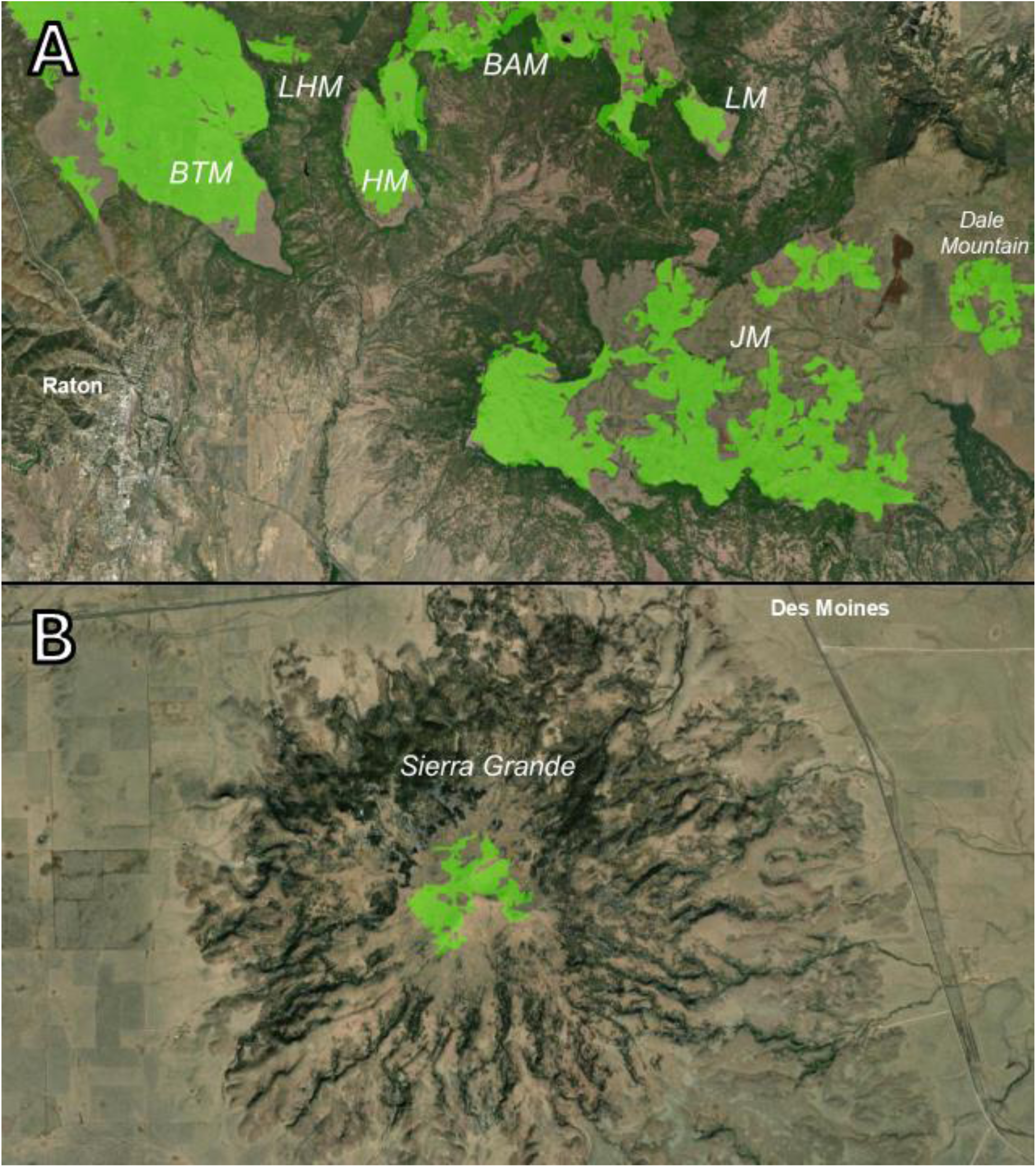
Results of habitat suitability model using minimum elevation of 2475m and minimum continuous area of 0.8km^2^. Basemap imagery by Esri et al. (2025). BTM = Barlett Mesa, HM = Horse Mesa, LHM = Little Horse Mesa, BAM = Barela Mesa, LM = Little Mesa, JM = Johnson Mesa.

## Discussion

In summary, from this investigation we were able to document new populations of Capulin arctics, verify the presence of historical populations, and assess population status at suspected sites of extirpation. Capulin arctics and their host plants were found at Johnson Mesa including in large numbers at Site 1. Our smaller numbers at Site 2 could be explained by visit times later in the afternoon. The butterfly and its host plants were also found on Bartlett Mesa. This is the first confirmation of the Capulin arctic on Bartlett Mesa. Much remains to be learned here, especially because the eastern edge is lower in elevation (2470–2530 m), with higher portions to the west and north attaining 2620 m). Capulin arctics were found historically at Sierra Grande although, to our knowledge, the last sighting of the Capulin arctic there was 17 May 1998.

Populations of Capulin arctics were also present at Capulin Volcano National Monument (CAVO) historically (the subspecies was named after the site). However, no Capulin arctics have been documented at CAVO since 1989 (Johnson *et al*., 2004). These surveys at Sierra Grande and CAVO suggest that populations of Capulin arctics do not occur at those sites despite the presence of host plants and otherwise suitable habitat. Little Horse Mesa also used to have a population, but no butterflies and few host plants were found on this recent visit. Little Horse Mesa was one of the lowest elevation mesas ever known to have supported the Capulin arctic with the last known sighting occurring in 2007. The mesa top is very grassy, and fescue is included, but it seems to have been superseded by other grasses in most places. An additional problem may be that Ponderosa pine and other trees are very prominent on much of the mesa.

The 2011 Track Fire removed a few trees, and young pines appear to be invading the meadows.

Finally, we used our habitat observations and host plant verification to model suitable habitat from public remote sensing data. These models predicted suitable habitat in seven geographic regions in in New Mexico: Sierra Grande, Johnson Mesa, Little Mesa, Barela Mesa, Horse Mesa, Little Horse Mesa, and Barlett Mesa. The mesas in Colfax County are near one another and those in Colorado such that they could fall within a reasonable dispersal distance for an arctic butterfly. However, the only habitat identified within Union County is an area of 2.6 km^2^ on Sierra Grande, 42.8 km away from the nearest habitat on Johnson Mesa. Capulin arctic on Sierra Grande, if it persists, appears to represent a very isolated population. The majority of estimated suitable habitat from all models is on privately-owned land (82-84.5%) (Fig. 5).

**Figure 5.**
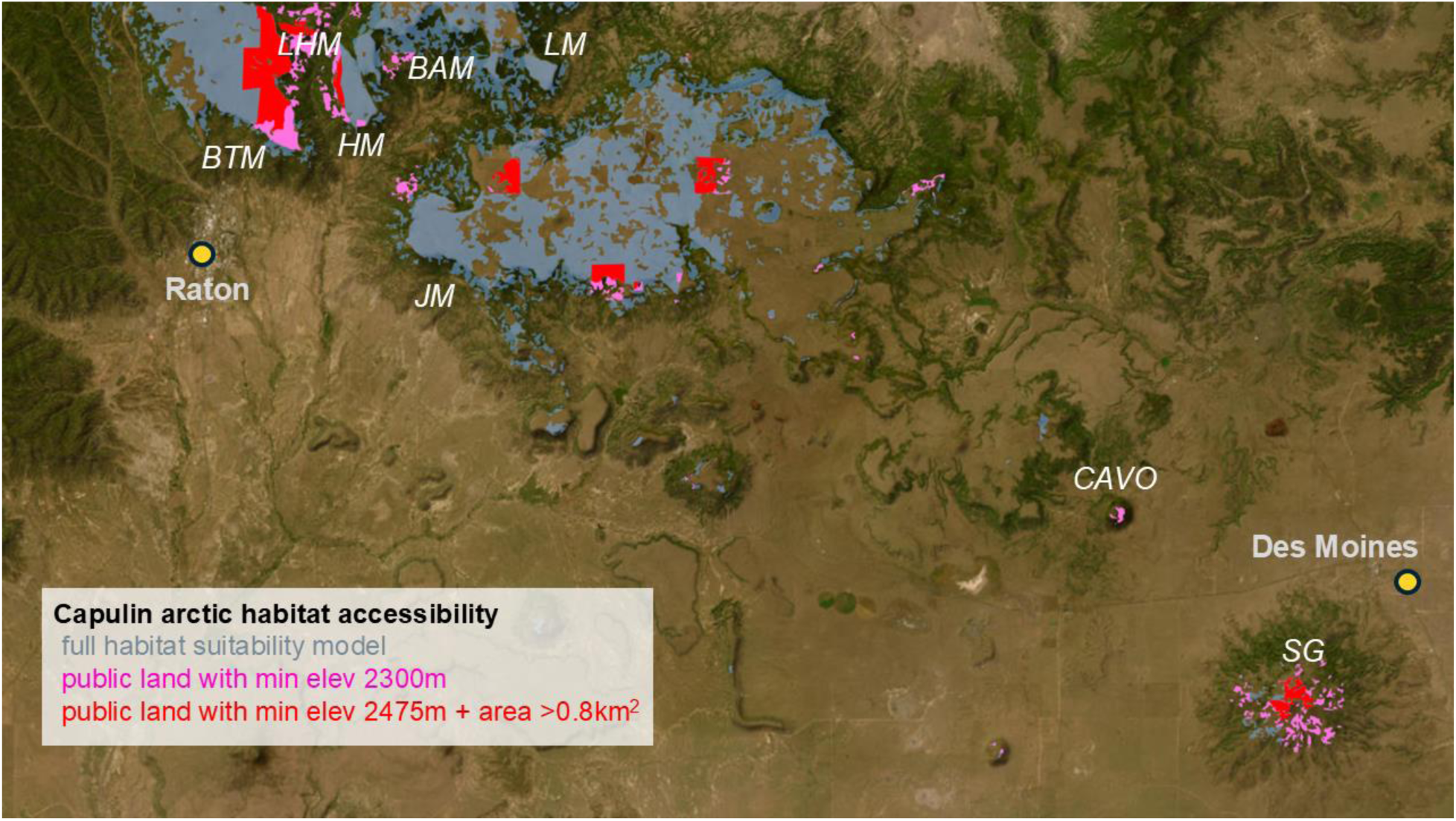
Capulin arctic potential habitat accessibility for future survey. Basemap imagery by Esri et al. (2025). BTM = Barlett Mesa, HM = Horse Mesa, LHM = Little Horse Mesa, BAM = Barela Mesa, LM = Little Mesa, JM = Johnson Mesa, CAVO = Capulin Volcano National Monument, SG = Sierra Grande.

Besides CAVO, the remaining public land parcels are managed by the State of New Mexico.

Modeling suitable habitat using remote sensing data, guided by hypotheses derived from natural history observations, has enabled more effective prediction of potential occurrences of this rare butterfly in future surveys. Analyses like this effectively produce a roadmap to improve our understanding of the distribution, populations, and ecological requirements of a threatened species, which is an essential step towards recovery.

This geographic data allowed us to calculate a well-supported estimate of the amount of suitable habitat available for the butterfly and can also guide future Capulin arctic investigations in New Mexico. This approach provides a fast and efficient way to estimate the potential maximum extent of the Capulin arctic’s range which can be ground-truthed over time. This may also provide a useful template for estimating the extent of suitable habitat for other arctic butterfly (*Oeneis)* species and subspecies, which is important as this genus is highly threatened in the southern United States. In New Mexico alone, four *Oeneis* are designated as SGCN.

Our mid-May field excursion seemed to hit the one-month adult flight period in its peak as we saw both males and females and both appeared freshly eclosed. Adults displayed expected behaviors, despite stiff midday and early afternoon winds. Most adults we saw were hidden in the grasses emerging only when we approached. When approached, the butterflies would take flight and continue for a couple meters before settling again. Others that took flight were caught by the wind and disappeared quickly from view. Their active flight rarely carried them more than a meter above ground level, or 2-3 m above *Festuca* stems. This observation that the butterfly rarely flies and when it does stays very close to the ground is consistent with observations at CAVO where it was originally recorded (Brown 1970). Brown (1970) also reported that he only observed males flying and all the females he saw were crawling or maneuvering on the ground. This may indicate that female Capulin arctics have an extremely short dispersal distance from their fescue host plants. Throughout the study area, nectar sources were few, and no nectaring was observed.

Our surveys revealed a close association between *Festuca idahoensis* and the Capulin arctic, strongly implicating this species as the butterfly’s primary larval host plant. Nevertheless, *F. idahoensis* represents only one component of the diverse grass community characteristic of the meadows and mesas where the butterfly occurs. In some sites, fescue was abundant and often among the dominant grasses, whereas in others it was sparse or apparently absent. During adult monitoring forays, Capulin arctic individuals were consistently most abundant in areas with high fescue cover and rare or absent in areas lacking fescue. These observations suggest that suitable habitat may be highly restricted within the broader landscape, likely limited to microhabitats containing dense stands or patches of *F. idahoensis*.

One particularly noteworthy case is Sierra Grande, where we observed extensive *Festuca* cover but no apparent evidence of the historically present Capulin arctic population. There are several possible explanations for this apparent local extirpation. At other sites where Capulin arctics persist, populations are restricted to flat, windswept, high-elevation mesas. In contrast, Sierra Grande is a singular volcanic peak, and the fescue-dominated habitat occurs on very steep upper slopes that transition downslope into a compact, flat basin at 2,410 m. If these steep slopes are unsuitable for the species, targeted resurveys of the basin may be warranted. Complicating matters, a perennial water source within the basin attracts cattle, possibly resulting in concentrated grazing pressure within the fescue meadow. It’s possible that sustained grazing may have degraded or eliminated critical butterfly habitat here. However, focused, site-specific research will be needed to determine if Capulin arctic is absent and, if it is to assess what factors may have driven the disappearance of the Sierra Grande colony.

Sierra Grande though is just one of three sites (Sierra Grande, CAVO, and Little Horse Mesa) where Capulin arctic colonies seem to have been extirpated. With the butterfly’s recent designation as a Species of Greatest Conservation Need, it is worth considering if natural recolonization from another mesa is possible. From observations made herein and in Brown (1970), the Capulin arctic is not a particularly strong flier or disperser compared with other butterflies, and this might limit its ability to colonize islands of suitable habitat on its own.

Additional support for limited dispersal ability comes from Brown’s (1970) observations at CAVO where he suggested that the butterfly might be experiencing inbreeding depression. However, on May 2, 2025, we observed a Capulin arctic in the lower topographic saddle between Bartlett Mesa and Little Horse Mesa, offering an example of a possible mechanism or "bridge" for dispersal between adjacent mesa habitats. We had previously documented one Capulin arctic at the eastern base of Bartlett Mesa on May 22, 2007, and at the time it was presumed that westerly winds had simply blown this individual off the top of Bartlett Mesa.

Given the recent observation of this same behavior, we now suspect this is a functional mechanism for inter-mesa dispersal for this insect. Therefore, re-colonization is theoretically possible, as long as the threats that resulted in extirpation do not persist.

It is also notable that CAVO (2493 m), Little Horse (2530 m) and the “bowl” at Sierra Grande (2377 m) have the lowest elevations of all known former and present colonies of the Capulin arctic. The Alberta arctic species in total is a cold-adapted boreal grassland specialist butterfly with most of its distribution in Canada and the northern United States (Cary & Toliver, 2025). Local extinction of the subspecies at the sites lowest in elevation may indicate that warming resulting from climate change has already affected this New Mexico subspecies. Many butterflies respond to warming average temperatures by moving to higher elevations or latitudes (Rödder *et al*., 2021). On the high elevation mesas and volcanic peaks in this area, populations are already inhabiting the highest elevations possible Further research is urgently needed to clarify these dynamics and to understand the broader impacts of climate change on the Capulin arctic, particularly for lowest-elevation populations, which may be the most vulnerable to future warming. This study not only supports the accurate and more complete conservation assessment of the Capulin arctic but also provides an example of the type of background research that is not only necessary, but achievable, to aid in the recovery of SGCN insect species.

With the ratification of New Mexico’s State Wildlife Action Plan in June 2025, the Capulin arctic is now designated a Species of Greatest Conservation Need. Having been extirpated from Capulin Volcano National Monument (CAVO) and potentially two other sites, the subspecies may require additional research and monitoring to ensure its persistence. We support establishing regular survey routes on state land where the butterfly remains extant to confirm occupancy and estimate population size, while continuing general butterfly monitoring at CAVO to detect possible recolonization and track other species inhabiting the Raton Mesa Complex. Further surveys should aim to confirm whether the Capulin arctic is truly extirpated from Sierra Grande, which appears beyond natural recolonization range, though potential reintroduction could be explored pending a clearer understanding of extirpation causes. To refine habitat models and assess meadow dynamics, it will be necessary to ground -truth *Festuca* distributions, evaluate entire plant communities, and monitor vegetation change using remote-sensing tools. Because two key sites are under the New Mexico State Land Office management, collaboration with that agency, as well as Colorado Parks and Wildlife, will be essential for coordinated research and recovery planning. Potential collaboration with private landowners in the region will further improve complete assessment of habitat. Finally, maintaining genetic health and connectivity among populations will require identifying potential dispersal corridors or, if necessary, considering assisted gene flow through targeted management interventions.

Lastly, it is important to recognize that the Capulin arctic is only one of 93 insects currently listed as SGCN in New Mexico, many of which face similarly urgent threats and remain poorly studied. The challenges confronting these species highlight a broader conservation crisis for the state’s insect fauna, emphasizing the need for systematic monitoring and data-driven management. We hope that the methodology presented here can serve as a practical framework, not only for the Capulin arctic, but also for other at-risk insect species, providing a foundation for ongoing efforts to document, monitor, and ultimately conserve New Mexico’s most vulnerable insect populations.

**Table 1.** Records of Capulin arctic butterfly in New Mexico. Updated from Johnson et al. (2004).

| <u>Location</u> | <u>Date</u> | <u>Observer/Source</u> |
| --- | --- | --- |
| Capulin Volcano NM, Union Co. | 22-April-1968 | G. Forbes |
| Capulin Volcano NM, Union Co. | 17-May-1969 | F. M. Brown<br>(1971[1972]) |
| Capulin Volcano NM, Union Co. | 18-May-1969 | F. M. Brown<br>(1971[1972]) |
| Capulin Volcano NM, Union Co. | 30-May-1969 | F. M. Brown<br>(1971[1972]) |
| Capulin Volcano NM, Union Co. | 23-May-1970 | F. M. Brown<br>(1971[1972]) |
| Capulin Volcano NM, Union Co. | 30-May-1970 | M. E. Toliver |
| Capulin Volcano NM, Union Co. | 5-Jun-1971 | M. E. Toliver |
| Capulin Volcano NM, Union Co. | 3-May-1972 | J. Scott |
| Capulin Volcano NM, Union Co. | 20-May-1981 | F. & J. Preston |
| Capulin Volcano NM, Union Co. | 21-May-1981 | F. & J. Preston |
| Capulin Volcano NM, Union Co | 25-May-1989 | F. & J. Preston |
| Raton Mesa E of Raton Pass, Colfax Co. | 3-May-1972 | J. Scott |
| near Raton, Colfax Co. | 19-May-1986 | T. Kral |
| Sierra Grande, Union Co. | May-1993 | M. Fisher |
| Sierra Grande, Union Co. | 3-May-1997 | S. J. Cary |
| Sierra Grande, Union Co. | 31-May-1997 | S. J. Cary |
| Sierra Grande, Union Co. | 17-May-1998 | S. J. Cary |
| Johnson Mesa, Colfax Co. | 20-May-1990 | F. & J. Preston |
| Johnson Mesa, Colfax Co. | 12-May-1996 | S. J. Cary & R. Holland |
| Johnson Mesa, Colfax Co. | 12-May-2000 | S. J. Cary |
| Little Horse Mesa, Sugarite Canyon State Park,<br>Colfax Co. | 12-May-2000 | S. J. Cary |
| Johnson Mesa, Colfax Co. | 27-May-2003 | S. J. Cary & K. Johnson |
| Johnson Mesa, Colfax Co. | 20-May-2004 | S. J. Cary & K. Johnson |
| Little Horse Mesa, Sugarite Canyon State Park,<br>Colfax Co. | 21-May-2004 | S. J. Cary |
| saddle between Bartlett and Little Horse mesas | 22-May-2007 | S. J. Cary |
| Horse Mesa, Sugarite Canyon State Park, Colfax Co. | 20-May-2008 | S. J. Cary |
| Johnson Mesa, Colfax Co. T31N R25E S16 | 13-May-2025 | S. J. Cary |
| Johnson Mesa, Colfax Co. T31N R26E S16 | 14-May-2025 | S. J. Cary & Q. Baine |
| Johnson Mesa, Colfax Co. | 15-May-2025 | S. J. Cary, Q. Baine, S. Doneski, K. Ricks |
| Bartlett Mesa, Colfax Co., NM | 16-May-2025 | S. J. Cary, S. Doneski, Q. Baine |
| saddle between Bartlett and Little Horse mesas | 24-May-2025 | K. Ricks |

## Acknowledgements

The authors are grateful to Anna Walker for managing the New Mexico Butterfly Monitoring Network; David Lightfoot for access to lab equipment; Lysandra Pyle and Peri Lee Pipkin for grass identification support; Jennifer Pedneau for field survey support; and Matt Forister for suggestions to improve this manuscript. Land access was granted by the U.S. Fish and Wildlife Service to Capulin Volcano National Monument (Scientific Collecting and Research Permit CAVO-2025-SCI-0001), and by the New Mexico State Land Office to state trust lands (Natural Resource Authorization #FOD-NR-440 and #RA-2025-03114).

